## supplemental files for "Bidirectional crosstalk between Hypoxia-Inducible Factor and glucocorticoid signalling in zebrafish larvae"

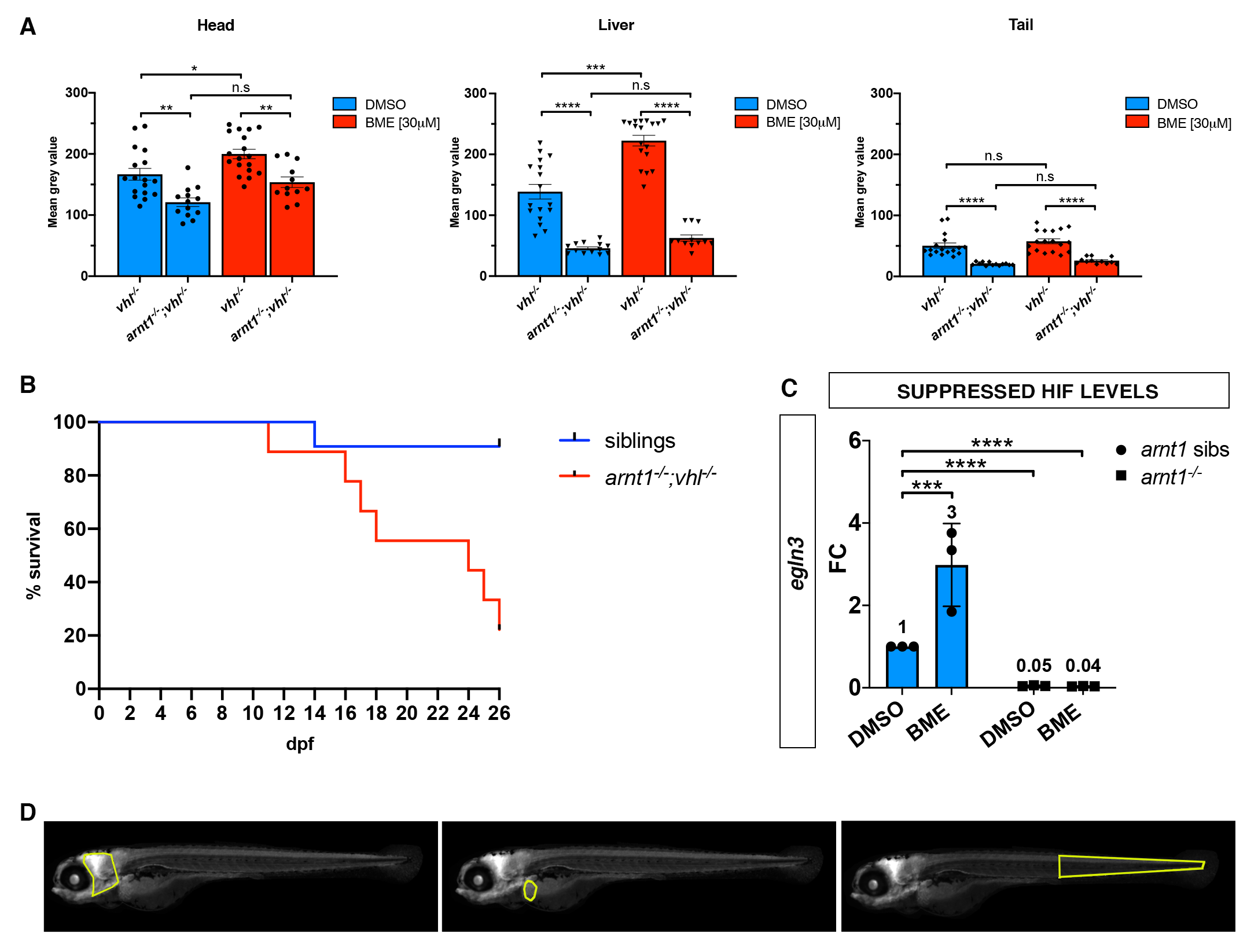
**Supporting information**

**Fig S1. arnt1^-/-^; vhl^-/-^ larvae showed a reduced phd3:eGFP brightness and a partially rescued vhl phenotype.**

A. Statistical analysis performed on mean gray value quantification (at the level of the head, liver and tail), after phenotypic analysis on 5dpf DMSO and BME [30μM] treated arnt1^+/-^;vhl^+/-^(phd3:eGFP) x arnt1^-/-^; vhl^+/-^(phd3:eGFP) derived larvae (n=540). vhl^-/-^ DMSO treated n=17 larvae: head 166.67 ± 9.63 (mean ± s.e.m); liver 138.61 ± 12.05 (mean ± s.e.m); tail 50.31 ± 4.51 (mean ± s.e.m). arnt1^-/-^;vhl^-/-^ DMSO treated n = 13 larvae: head 121.05 ± 6.99 (mean ± s.e.m); liver 49.61 ± 3.88 (mean ± s.e.m); tail 21.75 ± 1.12 (mean ± s.e.m). vhl^-/-^ BME treated n = 18 larvae: head 199.88 ± 7.71 (mean ± s.e.m); liver 222.57 ± 8.72 (mean ± s.e.m); tail 57.57 ± 4.11 (mean ± s.e.m). arnt1^-/-^;vhl^-/-^ BME treated n = 12 larvae: head 153.71 ± 8.66 (mean ± s.e.m); liver 62.58 ± 5.16 (mean ± s.e.m); tail 25.82 ± 1.54 (mean ± s.e.m). Ordinary One-way ANOVA followed by Sidak’s multiple comparison test (*P < 0.05; **P < 0.01; ***P <0.001; ****P < 0.0001).

B. Kaplan-Meier survival curves of the zebrafish arnt1^+/-^; vhl^+/-^(phd3:eGFP) genotype analysed in this study. Time is shown in days. Siblings n = 30; arnt1^-/-^; vhl^-/-^(phd3:eGFP) n = 8. The Log-rank (Mantel-Cox) test was used for statistical analysis. arnt1^-/-^; vhl^-/-^(phd3:eGFP) vs. siblings: **P < 0.0027.

C. RTqPCR analysis performed on arnt1 siblings (n=10; 3 repeats) and arnt1^-/-^ (n=10; 3 repeats) larvae at 5 dpf, using egln3 as probe. Statistical analysis was performed on ΔΔCt values, whereas data are shown as fold change values. Ordinary Two-way ANOVA followed by Dunnett’s multiple comparison test (***P <0.001;****P < 0.0001).

D. Representative picture of head, liver and tail areas selected in each larva to quantify the phd3:eGFP-related brightness via mean grey value quantification (Fiji, ImageJ software).

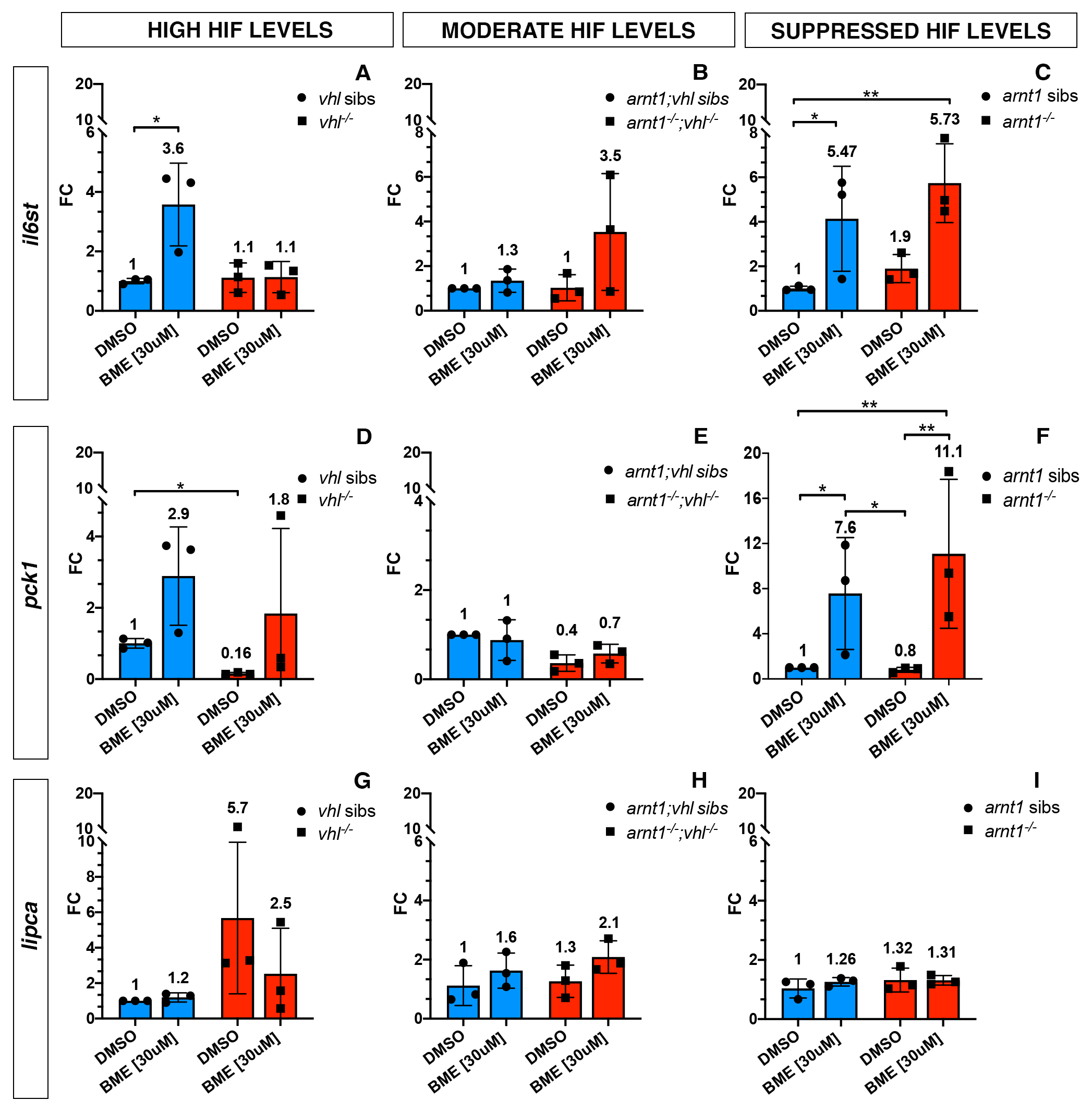

**Fig S2. GC target genes expression in the presence of high, moderately upregulated and suppressed HIF signalling pathway.**

Schematic view of RTqPCR analysis on il6st, pck1 and lipca (GC target genes) expression performed on the following mutant lines: vhl+/-(phd3:eGFP), arnt1+/-;vhl+/-(phd3:eGFP) and arnt1+/-(phd3:eGFP). Statistical analysis performed on Ct values; data are shown as fold change values for RTqPCR analysed samples; ordinary Two-way ANOVA followed by Dunnett’s multiple comparison test (*P < 0.05; **P < 0.01; ***P <0.001; ****P < 0.0001).

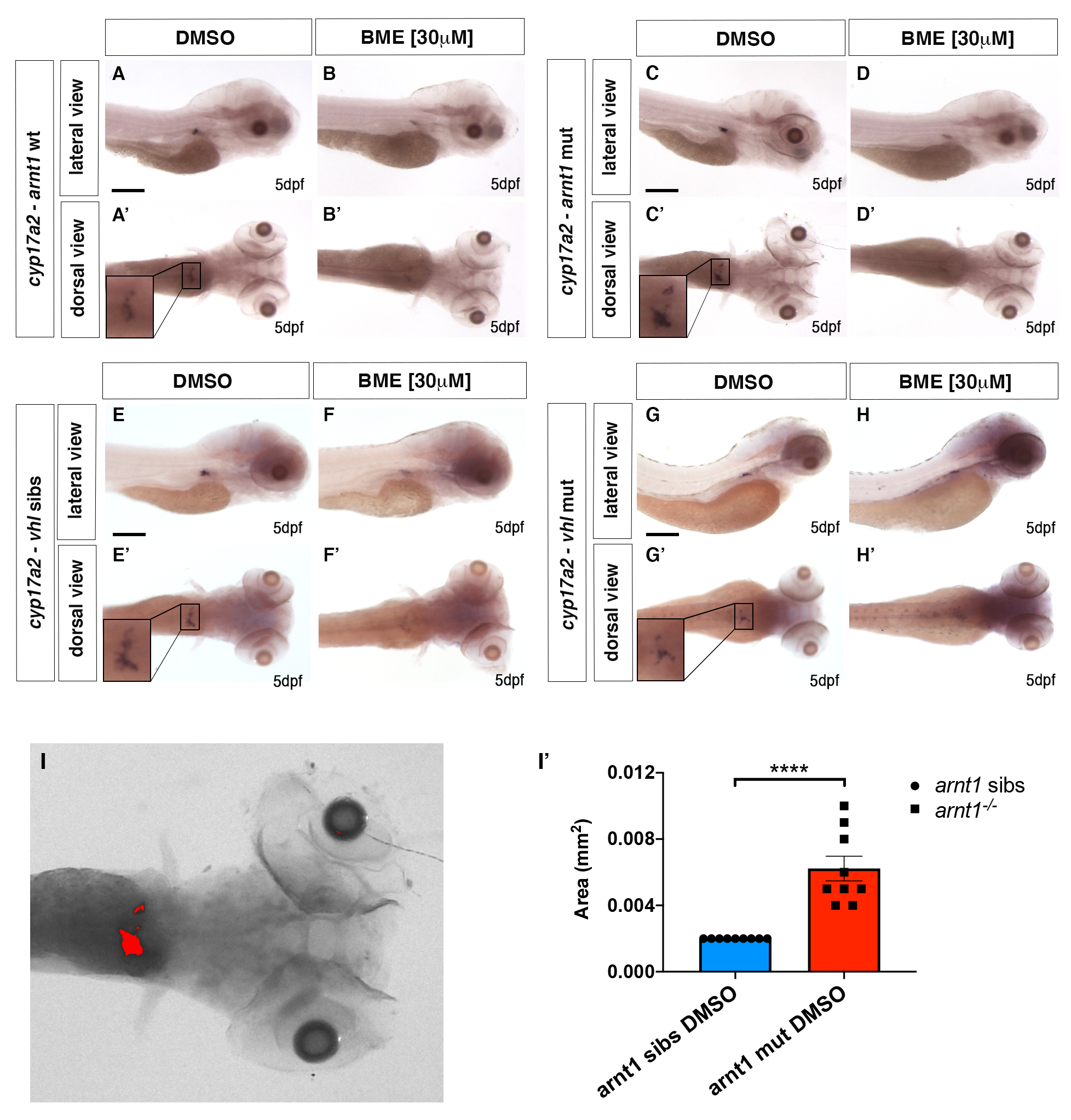

**Fig S3. gr^-/-^; vhl^-/-^ larvae showed a reduced phd3:eGFP brightness and a partially rescued Vhl phenotype.**

A-D’. Representative pictures of WISH performed on DMSO and BME [30 μM] treated *arnt1* mutant line, at 5 dpf, using *cyp17a2* as probe. A-A’) *arnt1* wt DMSO treated larvae (n= 26/28) showed normal *cyp17a2* expression, whereas 2/28 larvae showed a weaker one; B-B’) *arnt1* wt BME treated larvae (n= 28/30) showed downregulated *cyp17a2* expression, whereas 2/30 larvae showed a normal one. C-C’) In contrast, *arnt1*^-/-^ DMSO treated larvae (n= 24/28) showed upregulated *cyp17a2* expression, whereas 4/28 larvae showed a weaker *one.* D-D’) *arnt1*^-/-^ BME treated larvae (n= 25/29) showed downregulated *cyp17a2* expression, whereas 4/29, showed a normal one. Chi-square test (****P < 0.0001). Scale bar 200 μm.

E-H’. Representative pictures of WISH performed on DMSO and BME [30 μM] treated *vhl* mutant line, at 5 dpf, using *cyp17a2* as probe. E-E’) DMSO treated *vhl* siblings (n= 18/21) showed normal *cyp17a2* expression, whereas 3/21 larvae showed a weaker one; F-F’) BME treated *vhl* siblings (n= 28/30) showed downregulated *cyp17a2* expression, whereas 2/30 larvae showed a normal one. G-G’) On the other hand, *vhl*^-/-^ DMSO treated larvae (n= 27/28) showed weak *cyp17a2* expression, whereas 1/28 larvae showed a normal *one.* H-H’) *vhl*^-/-^ BME treated larvae (n= 30/30) showed downregulated *cyp17a2* expression. Chi-square test (****P < 0.0001). Scale bar 200 μm.

I-I’. Representative picture of the colour threshold area calculation method (ImageJ software’s tool) used to quantify the area occupied by the *cyp17a2* WISH staining both in *arnt1* siblings (n=9) and *arnt1*^-/-^ (n=9). I’. unpaired t-test (****P <0.0001).

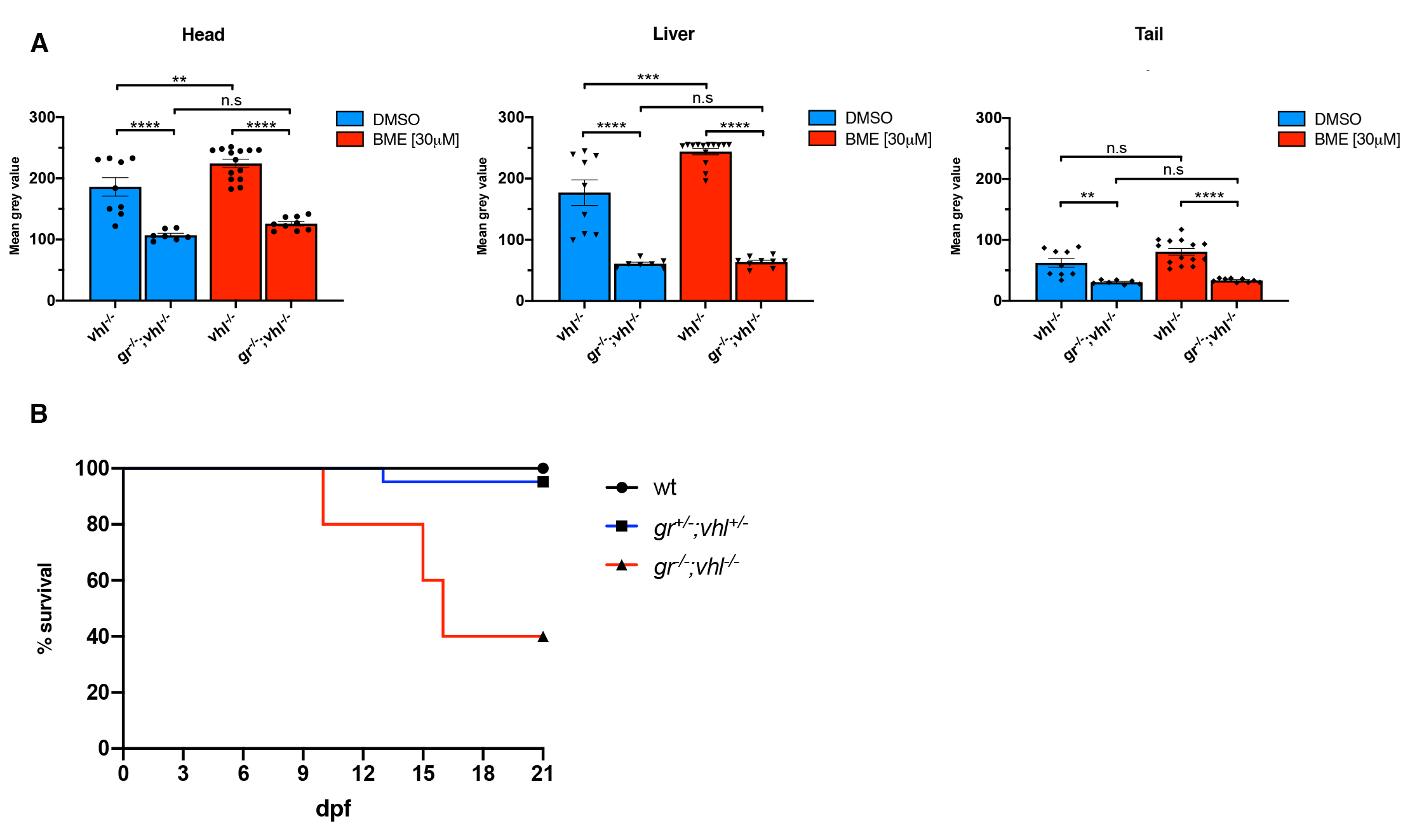

**Fig S4. gr^-/-^; vhl^-/-^ larvae showed a reduced phd3:eGFP brightness and a partially rescued vhl phenotype.**

A. Statistical analysis performed on mean gray value quantification (at the level of the head, liver and tail), after phenotypic analysis on 5dpf DMSO and BME [30μM] treated gr^+/-^;vhl^+/-^(phd3:eGFP) x gr^-/-^; vhl^+/-^(phd3:eGFP) derived larvae (n=600). vhl^-/-^ DMSO treated n = 9 larvae: head 186 ± 15.12 (mean ± s.e.m); liver 177.01 ± 20.85 (mean ± s.e.m); tail 62.34 ± 7.27 (mean ± s.e.m). gr^-/-^;vhl^-/-^ DMSO treated n = 7 larvae: head 106.96 ± 3.21 (mean ± s.e.m); liver 60.75 ± 2.56 (mean ± s.e.m); tail 30.67 ± 1.27 (mean ± s.e.m). vhl^-/-^ BME treated n = 14 larvae: head 224.32 ± 6.83 (mean ± s.e.m); liver 244.07 ± 5.31 (mean ± s.e.m); tail 80.51 ± 5.49 (mean ± s.e.m). gr^-/-^;vhl^-/-^ BME treated n = 9 larvae: head 125.85 ± 3.6 (mean ± s.e.m); liver 63.56 ± 2.91 (mean ± s.e.m); tail 33.67 ± 1.02 (mean ± s.e.m). Ordinary One-way ANOVA followed by Sidak’s multiple comparison test (*P < 0.05; **P < 0.01; ***P <0.001; ****P < 0.0001).

B. Kaplan-Meier survival curves of the zebrafish gr^+/-^; vhl^+/-^(phd3:eGFP) genotype analysed in this study. Time is shown in days. Wild-types n = 20; gr^+/-^; vhl^+/-^ n = 20; gr^-/-^; vhl^-/-^(phd3:eGFP) n = 5. The Log-rank (Mantel-Cox) test was used for statistical analysis. gr^-/-^; vhl^-/-^(phd3:eGFP) vs. gr^+/-^; vhl^+/-^, ****P < 0.0001; gr^-/-^; vhl^-/-^(phd3:eGFP) vs. wt, ****P < 0.0001.

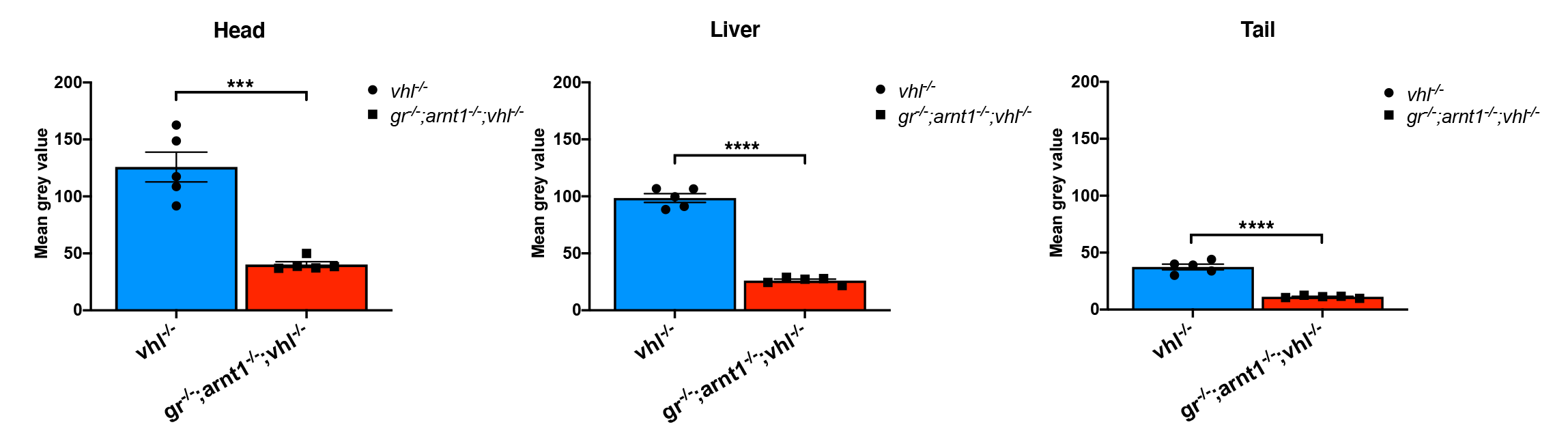

**Fig S5. gr^-/-^;arnt1^-/-^;vhl^-/-^ showed an even more reduced phd3:eGFP brightness.**

Statistical analysis performed on mean gray values quantification (at the level of the head, liver and tail), after phenotypic analysis on 5dpf gr^+/-^;arnt1^+/-^vhl^+/-^(phd3:eGFP) incross-derived GFP^+^ larvae (n=488). vhl^-/-^ n = 5 larvae: head 125.82 ± 13.05 (mean ± s.e.m); liver 98.52 ± 3.8 (mean ± s.e.m); tail 37.43 ± 2.45 (mean ± s.e.m). gr^-/-^;arnt1^-/-^;vhl^-/-^ n = 5 larvae: head 40.24 ± 2.46 (mean ± s.e.m); liver 26.07 ± 1.31 (mean ± s.e.m); tail 11.22 ± 0.47 (mean ± s.e.m); unpaired t-test (***P = 0.0002; ****P < 0.0001).

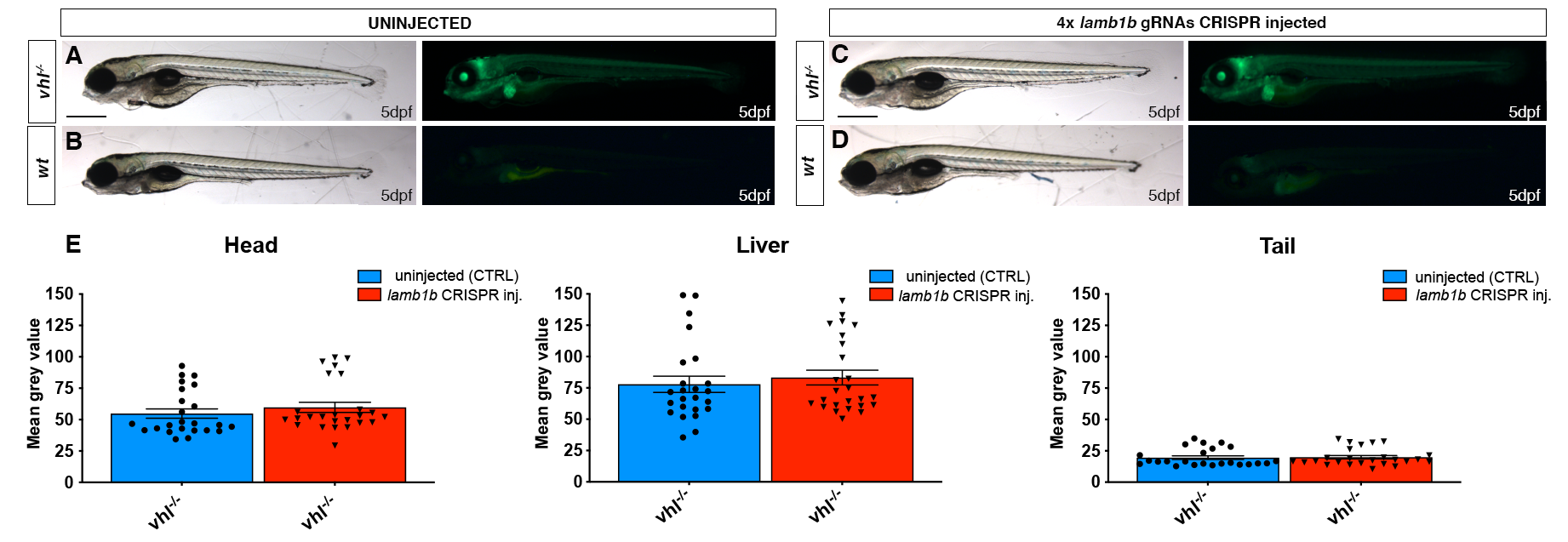

**Fig S6. CRISPR/Cas9 injection per se does not affect HIF signalling.**

A-D. Representative pictures of 5 dpf CRISPANT mutants created by redundantly targeting lamb1b gene via co-injection of 4x gRNAs in vhl^+/-^(phd3:eGFP) incross-derived embryos (n=400). Uninjected embryos were used as control (n=470). Fluorescence, exposure = 991,4 ms. Scale bar 500 μm.

E. Statistical analysis performed on mean grey values quantification (at the level of the head, liver and tail), after phenotypic analysis on 5 dpf lamb1b 4x gRNAs injected and uninjected vhl^+/-^(phd3:eGFP) incross-derived larvae. vhl^-/-^ uninjected n = 24 larvae: head 54.83 ± 3.68 (mean ± s.e.m); liver 77.86 ± 6.46 (mean ± s.e.m); tail 19.56 ± 1.43 (mean ± s.e.m). vhl^-/-^ injected n = 25 larvae: head 59.74 ± 4.05 (mean ± s.e.m); liver 83.23 ± 5.92 (mean ± s.e.m); tail 19.9 ± 1.38 (mean ± s.e.m); unpaired t-test (all panels: *P < 0.05; **P < 0.01; ***P <0.001; ****P < 0.0001).
